# Endpoint PCR coupled with capillary electrophoresis (celPCR) provides sensitive and quantitative measures of environmental DNA in singleplex and multiplex reactions

**DOI:** 10.1101/2020.10.24.353730

**Authors:** Bettina Thalinger, Yannick Pütz, Michael Traugott

**Author notes:** **Corresponding author:** Bettina Thalinger, Centre for Biodiversity Genomics, University of Guelph, 50 Stone Road E, N1G 2W1, Guelph, Ontario, Canada.

## Abstract

The use of sensitive methods is key for the detection of target taxa, from trace amounts of environmental DNA (eDNA) in a sample. In this context, digital PCR (dPCR) enables direct quantification and is commonly perceived as more sensitive than endpoint PCR. However, endpoint PCR coupled with capillary electrophoresis (celPCR) potentially embodies a viable alternative as it quantitatively measures signal strength in Relative Fluorescence Units (RFU). Provided comparable levels of sensitivity are reached, celPCR permits the development of cost-efficient multiplex PCRs, enabling the simultaneous detection of several target taxa.

Here, we compared the sensitivity of singleplex and multiplex celPCR to dPCR for species-specific primer pairs amplifying mitochondrial DNA (COI) of fish species occurring in European freshwaters by analysing dilution series of DNA extracts and field-collected water samples. Both singleplex and multiplex celPCR and dPCR displayed comparable sensitivity with reliable positive amplifications starting at two to 10 target DNA copies per µl DNA extract. celPCR was suitable for quantifying target DNA and direct inference of DNA concentrations from RFU was possible after accounting for primer effects. Furthermore, multiplex celPCRs and dPCRs were successfully used for the detection and quantification of fish-eDNA in field-collected water samples, confirming the results of the dilution series experiment and exemplifying the high sensitivity of the two approaches.

The possibility of detection and quantification via multiplex celPCR is appealing for the cost-efficient screening of high sample numbers. The present results confirm the sensitivity of this approach thus enabling its application for future eDNA-based monitoring efforts.

## Introduction

DNA traces contained in environmental samples are frequently used for the detection of species in environmental studies and wildlife biology [1]. Recently, species detection from water samples using environmental DNA (eDNA) - DNA fragments released in the form of excretions, secretions, and other bits of organisms into the environment [2] - has also moved from a purely scientific method to the successful application in routine species monitoring [3–7]. This creates a need for cost-efficient and reliable processing of large sample numbers.

Studies investigating the general species composition in environmental samples usually employ metabarcoding [6,8,9]. Individual species and their distribution are mainly investigated via targeted eDNA assays using endpoint PCR, quantitative real-time PCR (qPCR), or digital PCR (dPCR) [10–12]. For the amplification of eDNA, qPCRs and dPCRs are frequently complemented with probes to increase target-specific amplification. In addition, both techniques allow the quantification of target DNA [11,13]. Nevertheless, qPCR is an indirect approach as DNA quantities are calculated using standard curves and only dPCR enables direct and absolute DNA quantification [14]. Endpoint PCR is also commonly used to detect target DNA from environmental samples. Although the visualisation of amplification success on agarose gels and the resulting binary (yes/no) data can be used for occupancy modelling [15,16], it does not generally allow for quantitative estimates. This disadvantage can be compensated by analysing the endpoint PCR product via capillary electrophoresis (celPCR): in capillary electrophoresis all double-stranded DNA fragments are separated by their size and the amount of each fragment is quantified in a relative manner by measuring the Relative Fluorescence Units (RFU) of each fragment. This is possible as either the primers or the whole fragment is fluorescently labelled [17,18]. In the past, celPCR has been used to determine if the fluorescence of a target amplicon exceeds a predefined threshold and samples can thus be scored “positive” [19,20]. However, there has been only rudimentary attempts to assess the general quantification capabilities of celPCR for eDNA analyses [18,21]. This possibility for quantification is especially appealing for target eDNA detection in a large number of samples, as there is a high potential for cost-reduction based on PCR-chemicals alone (Table 1).

**Table 1:** Comparison of PCR reagent costs per reaction between commonly used kits for dPCR, qPCR and celPCR (prices in CAD are calculated from lot sizes of 5,000 reactions; retrieved on 24^th^ October 2020).

| PCR-type | supplier | product name | reaction volume [µl] | Price |
| --- | --- | --- | --- | --- |
| ceI PCR | Qiagen | Multiplex PCR Kit | 10 | 0.57 |
| dPCR | Bio-Rad | EvaGreen Supermix | 20 | 1.38 |
|  |  | Supermix for Probes | 20 | 1.38 |
| qPCR | Thermo Fisher Scientific | TaqPath™ qPCR Master Mix, CG | 20 | 1.57 |

Target DNA concentrations in environmental samples are usually low and therefore, the performance of both amplification and visualization methods at minute concentrations is crucial for the successful detection of target eDNA [22]. To compare the sensitivity of assays, the Limit of Detection (LOD) is commonly used, however, its definition differs between PCR platforms: For qPCR, it is frequently defined as the target DNA concentration at which 95% of the reactions yield a positive result [23,24]. Theoretically, dPCR requires three out of 3,000 droplets to be positive, albeit the detection of single molecules is considered viable [25]. In practice, the LOD was found to be below 0.5 copies per µl in the dPCR mix [22,26]. In celPCR, the objective quantification of the fluorescence signal enables the definition of an LOD, which so far was defined as the amount of target DNA copies from which a reliable positive amplification (i.e. three or more positive replicates) is possible [17,27]. Endpoint PCR is sometimes associated with reduced sensitivity in comparison to qPCR and dPCR [28,29]. However, the LODs determined for invertebrate and vertebrate DNA with celPCR (10 to 30 target DNA copies in the reaction [17,18,27]) are similar to qPCR LODs ranging from five to 50 copies in PCR [28,30,31]. celPCR can therefore be considered sufficiently sensitive for detecting minute eDNA quantities.

Another aspect of targeted DNA amplification, which is hardly used in combination with eDNA detection, is multiplexing, i.e. the amplification of more than one target DNA fragment via the simultaneous use of several taxon-specific primer pairs [17,32]. Independent of the PCR platform and primer specificity, multiplex PCRs need to be balanced to exhibit similar levels of sensitivity for each of the primer pairs used [17,33]. This can be achieved by designing primers with similar melting temperature while minimizing cross-reactivity and competition among them [17,34]. It is possible to adjust the concentration of specific primers or probes in PCR to counteract such effects [17]. In celPCR, multiplexing is accomplished by combining primer pairs yielding amplicons of different size [17,32]. However, such balanced multiplex celPCR assays [27] were so far not examined for any remaining effects of primer identity after the optimization process (e.g. via direct comparison with dPCR results). Multiplex celPCR has been employed for the efficient screening of large sample sets to study trophic interactions [20,35], but not yet for eDNA studies. Albeit distinction via fragment length differences is also possible for qPCR and dPCR [36,37], multiplexes on these instruments frequently employ specific dyes (attached to the respective probes) for each target [34,37]. The limited number of available dyes and their potential influence on primer/probe properties in addition to of all the above mentioned factors [17,38], make the development of endpoint PCR / celPCR multiplexes more feasible in comparison to qPCR and dPCR (but see [39] for a high-throughput qPCR approach). Generally, the use of multiplex PCRs enhances the cost- and time-effectiveness of any screening for specific target taxa [17,27,34], but there has been no in-depth assessment whether this is possible without forfeiting sensitivity and whether it is truly beneficial compared to singleplex endpoint PCRs, qPCRs, and dPCRs, which are most commonly applied for the detection of individual taxa from environmental samples.

We designed species-specific primers for the mitochondrial cytochrome *c* oxidase subunit I (COI) gene of seven freshwater fish species occurring in Central Europe and optimized amplification conditions for singleplex celPCR, dPCR, and two multiplex celPCRs. The sensitivity was compared between the three approaches via a dilution series experiment, which also evaluated the potential to quantify target eDNA from celPCR results. Finally, field-collected water samples were analyzed with multiplex celPCR and dPCR with the aim of estimating target eDNA copy number. We hypothesize that H1) it is possible to estimate target DNA copy number from RFU obtained by celPCR, H2) primer identity affects PCR efficiency even if primer characteristics are chosen for maximum similarity between primer pairs, and H3) both singleplex and multiplex celPCR show sufficient sensitivity to detect and quantify eDNA of all target species in field samples.

## Materials and Methods

All laboratory work was carried out in a clean-room laboratory at the University of Innsbruck, equipped with an ultraclean overpressure air system, separate rooms for DNA extraction, PCR preparation, PCR execution and post-PCR work, always using laminar flow workbenches, DNA-free gloves and protective clothing. All surfaces were cleaned with 10% bleach and 70% ethanol prior to laboratory work and all workbenches were daily radiated with UVC-light for three hours.

### Primer design and PCR optimization

Species-specific primers were designed for seven fish species commonly occurring in rhithral freshwaters in Central Europe, namely *Cottus gobio*, *Oncorhynchus mykiss*, *Salvelinus fontinalis*, *Salvelinus umbla*, *Salmo trutta*, *Squalius cephalus*, and *Thymallus thymallus*. For this task, a custom reference sequence database containing the COI sequences of all Central European freshwater fish species was used [27]. Suitable priming regions were identified using BioEdit Version 7.3.5 [40] before using Primer Premier 5 (PREMIER Biosoft International) to design species-specific primer pairs with melting temperatures as close as possible to 60 °C, amplicon lengths between 89 and 226 bp, and minimizing potential formation of dimers and secondary structures. After initial singleplex PCR testing, primer pairs were arranged in two multiplex PCR assays with at least 20 bp length difference between amplicons, enabling target identification based on amplicon length in capillary electrophoresis. Multiplex PCR conditions were optimized and primer concentrations adjusted to obtain similar sensitivity and amplification efficiency across all primer pairs using standardized DNA templates [17,18,27]. The final singleplex and multiplex PCRs underwent specificity testing using muscle tissue extracts from Central European fish species focusing on the seven target fish species, closely related species, and species with only a small number of mismatches at the respective priming sites. Two to three extracts were used per species (see SI1 for an alignment of target species, non-target species, and primers). Primers were found to be species-specific and no non-target amplification occurred with the below reported PCR conditions.

Both singleplex and multiplex endpoint PCR assays were based on the Multiplex PCR Kit (Qiagen) and contained bovine serum albumin (BSA) and tetramethylammonium chloride (TMAC) to reduce inhibition and enhance specificity [41,42]. Each 10 µl reaction contained 1 × reaction mix, 5 µg BSA, 30 mM TMAC, the respective primer combinations (Table 2) and 3.2 µl extract. For the dilution series experiment, the master mix was altered by using only 1 µl extract (or its respective dilution) and adding 2.2 µl molecular grade water. The thermocycling conditions with optimum sensitivity and specificity on a Mastercycler^®^ nexus (Eppendorf) were 15 min at 95 °C, 35 cycles of 94 °C for 30 s, 65 °C for 3 min and 72 °C for 60 s and final elongation at 72 °C for 10 min. For amplicon separation and visualization after endpoint PCR, the capillary electrophoresis system QIAxcel Advanced and the software QIAxcel ScreenGel (version 1.4.0, Qiagen) with the method AM320 and 30 s injection time were used. If PCR products of the expected fragment length reached a signal strength ≥ 0.08 RFU, they were deemed positive and their RFUs were recorded. The singleplex and multiplex celPCRs were run in 96-well plates and contained at least two negative and two positive controls (approx. 100 target DNA copies per target species and reaction). All negative controls resulted negative; all positive controls delivered the expected target amplicon(s). Albeit the *Salvelinus umbla* primer pair was included in one of the optimized multiplex reactions, it was not used in any of the consecutive processes (i.e. optimization on the dPCR platform, dilution series experiment) and the species was never detected in field-collected samples.

**Table 2:**
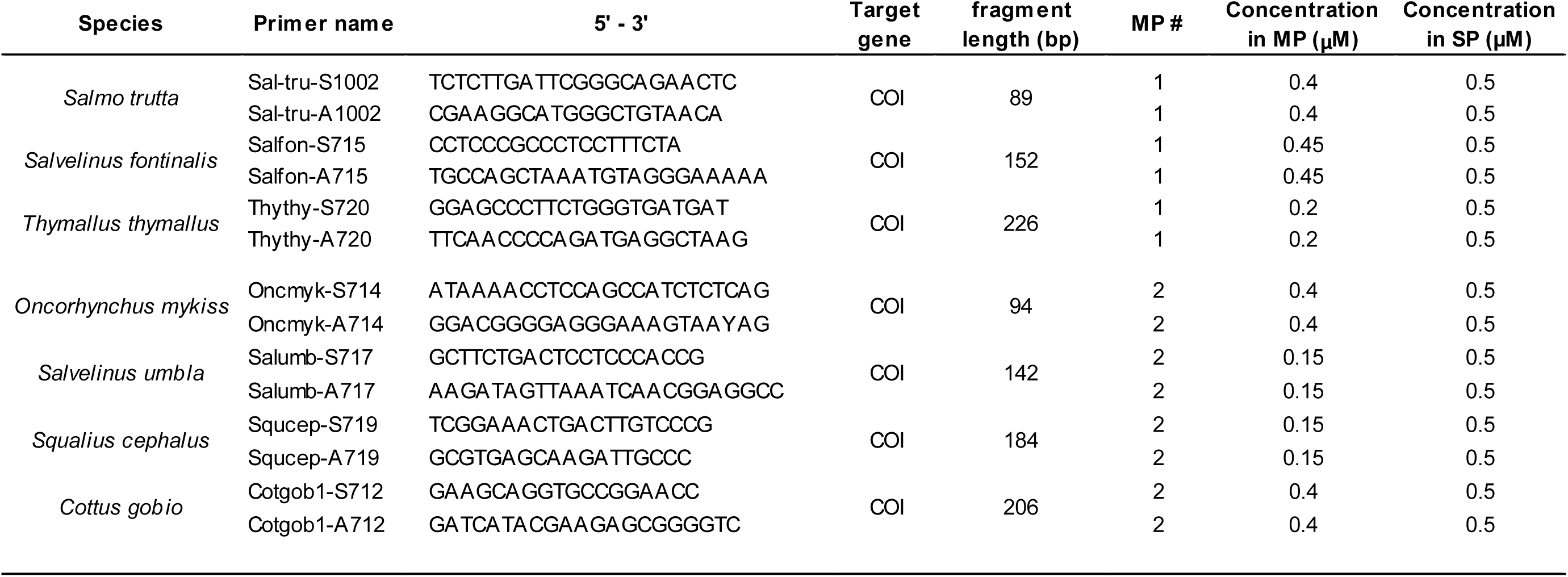
The target fish species and the associated species-specific primer pairs. The target gene, fragment length, association with one of the two multiplex assays (MP) and the respective primer concentrations in multiplex and singleplex celPCR are provided.

In a next step, the primer pairs (Table 2; exception: *S. umbla*) were used to create EvaGreen-based droplet dPCR assays using the AutoDG (Bio-Rad) for droplet generation, a Mastercycle^r®^ nexus for DNA amplification, and the QX200 Droplet Reader with its corresponding software QuantaSoft 1.0.596. (Bio-Rad) for fluorescence detection. We optimized PCR conditions by adjusting annealing temperature and/or time, and by using three-step protocols with separated annealing and extension phases to obtain a clear separation of positive and negative droplets and minimum “rain” (i.e. droplets with intermittent fluorescence between positive and negative droplets). Subsequently, a non-target test was conducted using the respectively other species and the three Central European fish species with the least sequence divergence at the priming sites. Ultimately, each 22 µl reaction mix, of which approx. 20 µl were used in the droplet generation process, contained 1 × EvaGreen Supermix (Bio-Rad) and 113.6 nM of each primer (Table 2) leaving 10.5 µl reaction volume, which was filled with 8.3 µl molecular grade water and 2.2 µl extract in the dilution series experiment, and varying extract volumes for the testing of field-collected samples. The optimum dPCR thermocycling conditions were 95 °C for 15 min, 40 cycles of 95 °C for 30 s, 58 °C (*O. mykiss* and *S. fontinalis*), or 60 °C (*S. trutta* and *S. cephalus*), or 62 °C (*T. thymallus*), or 64 °C (*C. gobio*) for 60 s, and 72 °C for 60 s, followed by stabilization at 4 °C for 5 min, 90 °C for 5 min, and 12 °C until further processing on the droplet reader. It was necessary to manually set a threshold for positive droplets for each target species, as the fluorescence levels varied with the fragment length generated by the respective primer pair. For *C. gobio* the threshold was set at 20,200 amplitude, for *O. mykiss* at 13,100, for *S. fontinalis* at 15,000, for *S. trutta* at 16,100, for *S. cephalus* at 18,300 and for *T. thymallus* at 18,400. All samples were processed in 96-well plates along with at least two positive and two negative controls, all of which resulted positive or negative, as expected.

### Dilution series experiment

The template DNA concentration of one extract from each of *C. gobio*, *O. mykiss, S. fontinalis*, *S. trutta*, *S. cephalus*, and *T. thymallus* was measured three times with the respective dPCR conditions described above. Based on these results, the extracts were diluted to 5,000 target DNA copies per µl extract using 1 × TE buffer. From there, a defined dilution series with 21 dilution steps (5,000; 4,000; 3,000; 2,000; 1,500; 1,000; 750; 500; 400; 300; 200; 150; 100; 80; 60; 40; 30; 20; 10; 5; 1 copy per µl) was generated. Each of the dilutions was used nine times: for three replicates of singleplex celPCR, multiplex celPCR, and dPCR under the conditions described above. For each species, the PCRs and the visualization of the obtained results were carried out right after setting up the dilution series. Cooling racks were used for each dilution and PCR preparation; diluted extracts were not frozen during processing. Throughout the experiment, each dPCR reaction produced more than 15,600 droplets (total) and the resulting concentrations were converted into target copies per µl for the respective dilution of the extract.

### Field samples

Per target species, 26 to 29 water samples, which were filtered and extracted as part of a larger field study (in prep.) were analyzed. For each sample, 2 L of water from different rivers in Tyrol (Austria) were collected in DNA-free wide-neck bottles and filtered in the field through 47 mm glass fibre filters with 1.2 µm mesh width (Whatman GF/C) using a peristaltic pump (Solinst, Model 410). Filters were transported in cooling boxes to the University of Innsbruck and stored at −20 °C until further processing. Cell lysis and DNA extraction were carried out as described by Thalinger et al. [18]: the filters were incubated overnight in lysis buffer before separating the extracts from the filters by centrifugation and extracting the DNA using the Biosprint 96 robotic platform (Qiagen).

All field samples were analyzed using the two multiplex PCR assays (Table 2) and capillary electrophoresis. For each of the species, 25 samples testing positive and five samples testing negative in multiplex celPCR were selected and analyzed with dPCR using the optimized conditions described above. To avoid background fluorescence from non-target DNA contained in the field sample extracts, 2.63 µl of extract was used per dPCR reaction for samples with RFUs above 0.5, 5.25 µl were used for samples with RFUs between 0.21 and 0.5, and 10.5 µl of extract was analyzed in case of RFUs between 0.08 and 0.2 to ensure a positive amplification despite very low target DNA concentration. As background fluorescence varied between samples from different locations, it was necessary to manually adjust the fluorescence threshold for positive droplets, albeit the positive and negative droplet clouds were clearly distinguishable for all samples.

### Statistical analysis

All calculations and visualizations were made in R Version 4.0.2 [43] using the packages “ggplot2”[44], “ggpubr”[45], “outliers”[46], “lme4”[47], “nlme”[48], and “MuMIn”[49].

First, the obtained RFUs and copy numbers from the singleplex celPCR, multiplex celPCR and dPCR were plotted against the expected copy numbers of the dilution series. Limits of Detection (LODs, i.e. the lowest number of target copies for which positive amplifications occurred; inferred from triplicate dPCR measurement of the same extract dilution) and Limits of Quantification (LOQs, i.e. all three replicates lead to a positive amplification) were evaluated for singleplex and multiplex celPCRs following Agersnap et al. [50] as it was not possible to directly transfer the LOD definition recently established by Klymus et al. [23] to this experiment. Prior to any other analyses, Grubbs’ tests were performed to remove outliers from the triplicate measurements [51]. Additionally, the lowest dilution was removed from the dataset, as not all replicates tested positive on all PCR platforms. Per dilution step and PCR method, the means and standard deviations of RFU and copies per µl extract were calculated. Based on these means, PCR efficiency was compared between RFU obtained from singleplex and multiplex celPCR using linear models. Then, the relationship between RFU and copies per µl extract was evaluated using linear mixed effects models. The natural logarithm of mean copies per µl extract was entered as independent variable, while mean RFUs derived from either singleplex or multiplex celPCR were entered as fixed effect, and fish species as random effect (random slope and intercept). As a next step, the models were used to predict copy number per µl from individual signal strengths for both singleplex and multiplex PCR results. Observed and predicted copy numbers were plotted against each other and for each species, a linear model and its 95% Confidence intervals (CI) were calculated. These models were compared to a 45 °-line representing the expected relation between observed and fitted copy numbers. Finally, linear models describing the relationship between *ln*-transformed copies and RFU in field samples were calculated, and observed and predicted copy numbers were plotted together with data obtained from the dilution series experiment.

## Results

In the dilution series experiment, the target DNA concentration per µl extract was quantified via dPCR for each of the six target species from a maximum of 23,680 copies to a minimum of 0.6 copies. Diluted extracts tested positive for all species with both singleplex and multiplex celPCRs, with RFU ranging from 0.09 to 6.53 in singleplex celPCR and 0.09 to 6.48 in multiplex celPCR, respectively. RFU showed an exponential decline with increasing dilution, and generally higher levels of variability (especially at higher DNA concentrations) compared to dPCR (Fig. 1).

**Figure 1:**
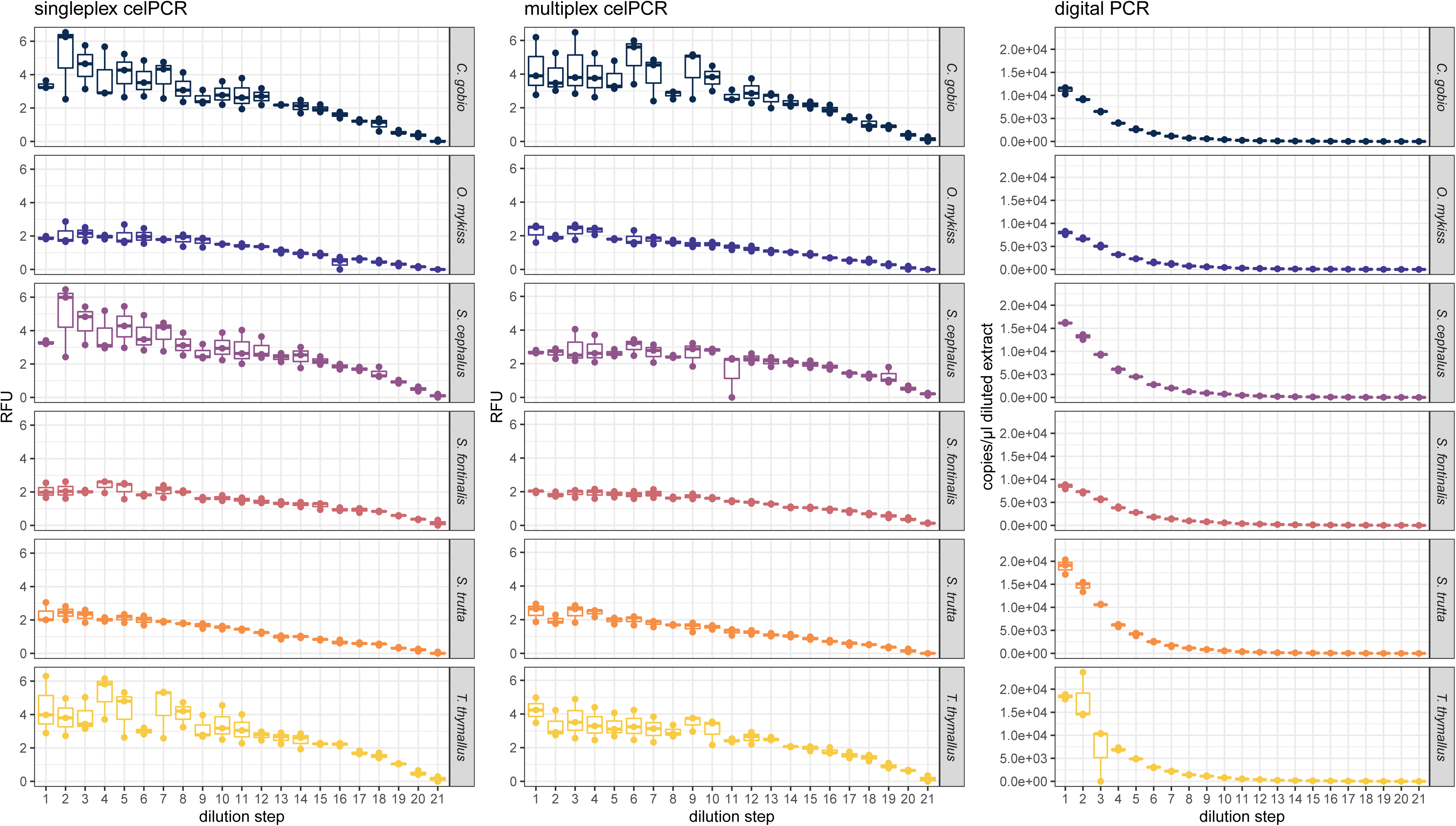
Relative Fluorescence Units (RFU) and template DNA copy numbers per µl diluted extract obtained for *C. gobio*, *O. mykiss, S. fontinalis*, *S. trutta*, *S. cephalus*, and *T. thymallus* from singleplex celPCR, multiplex celPCR, and dPCR. Dilution steps from 5,000 copies to 1 copy per µl extract are abbreviated 1 to 21.

Amplification efficiency differed significantly between singleplex and multiplex celPCRs for *S. cephalus*, *S. fontinalis*, and *T. thymallus*, with multiplex reactions leading to higher signal strengths at low DNA concentrations and singleplex reactions resulting in elevated RFU at high DNA concentrations (Fig. 2, SI2a). This trend was not observed for the three other species. The comparison of RFU (singleplex or multiplex celPCR) to copy numbery per µl extract obtained from dPCR showed amplification differences between primer pairs in endpoint PCR (Fig. 3a). After accounting for primer pair identity, *ln*-transformed copy number per µl extract could be predicted from singleplex and multiplex RFU (R² = 0.96 for both linear mixed effects models; Table 3, Fig. 3b). In both the singleplex and the multiplex celPCRs, the RFU produced by *C. gobio*, *T. thymallus* and *S. cephalus* primers were above the population mean (Fig. 3).

**Table 3:**
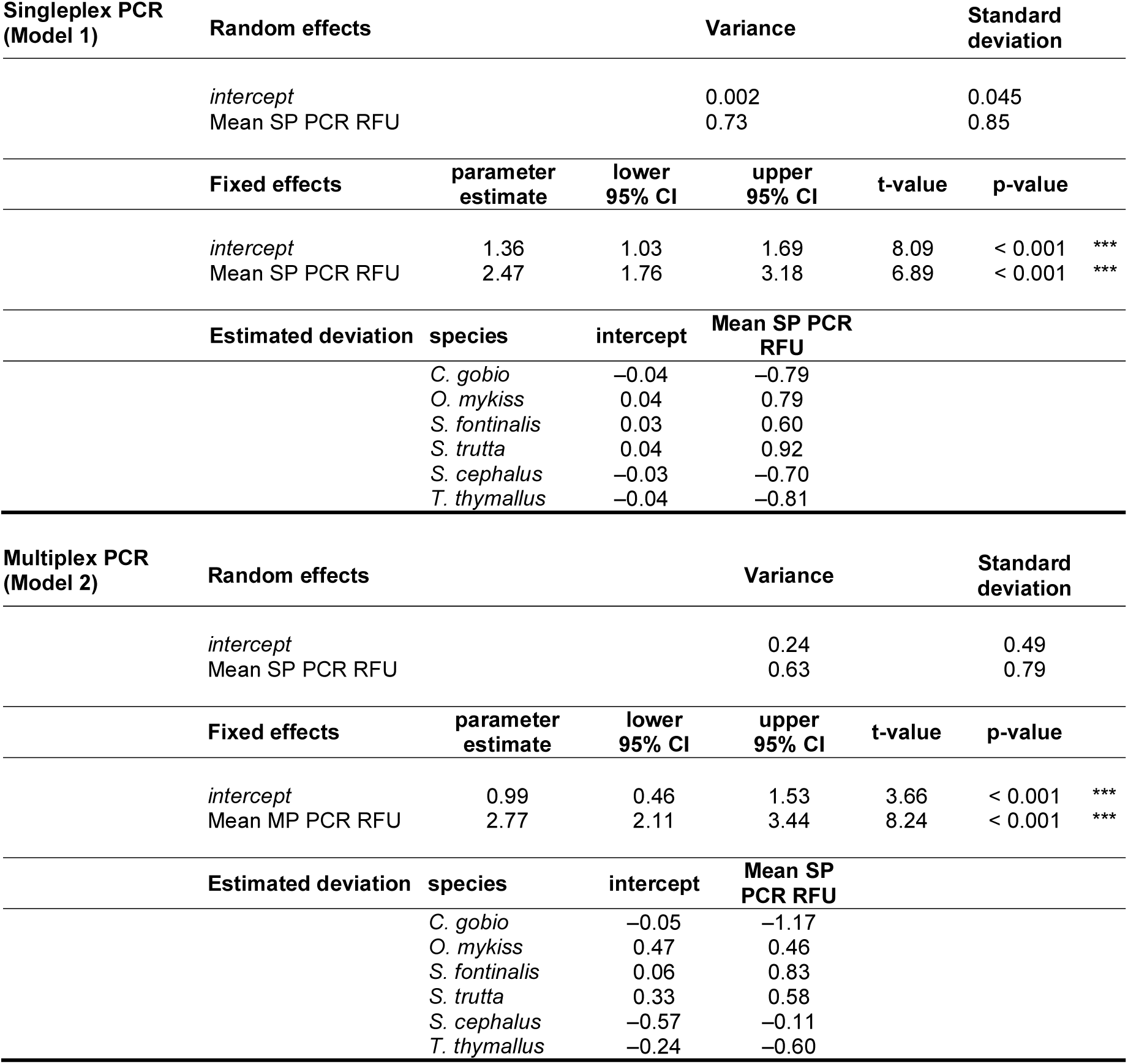
Linear mixed models for singleplex and multiplex celPCR

**Figure 2:**
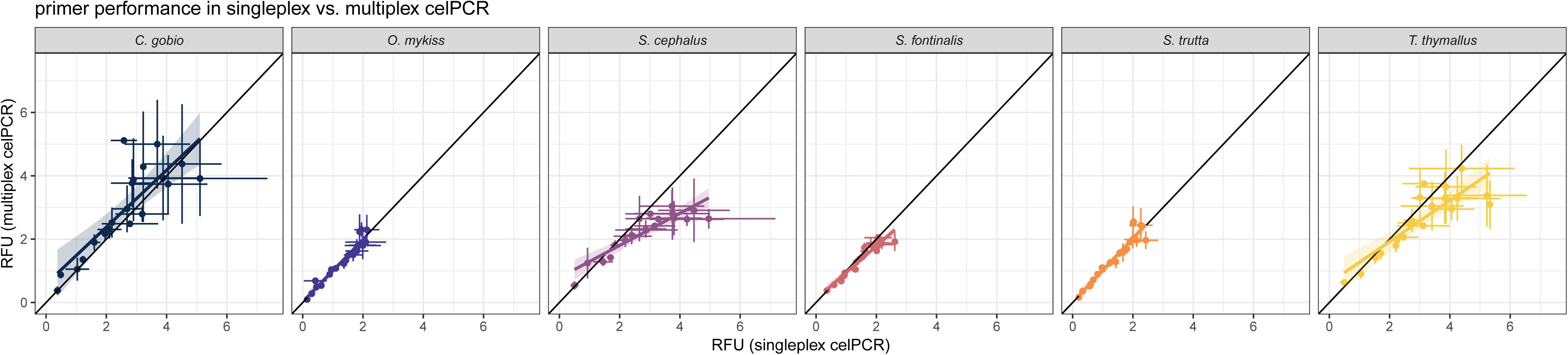
Primer performance comparison based on mean RFU (dots) obtained from singleplex and multiplex celPCRs. The corresponding standard deviations are displayed as whiskers; the shaded area depicts the 95%-CIs; see SI2 for model specifications.

**Figure 3:**
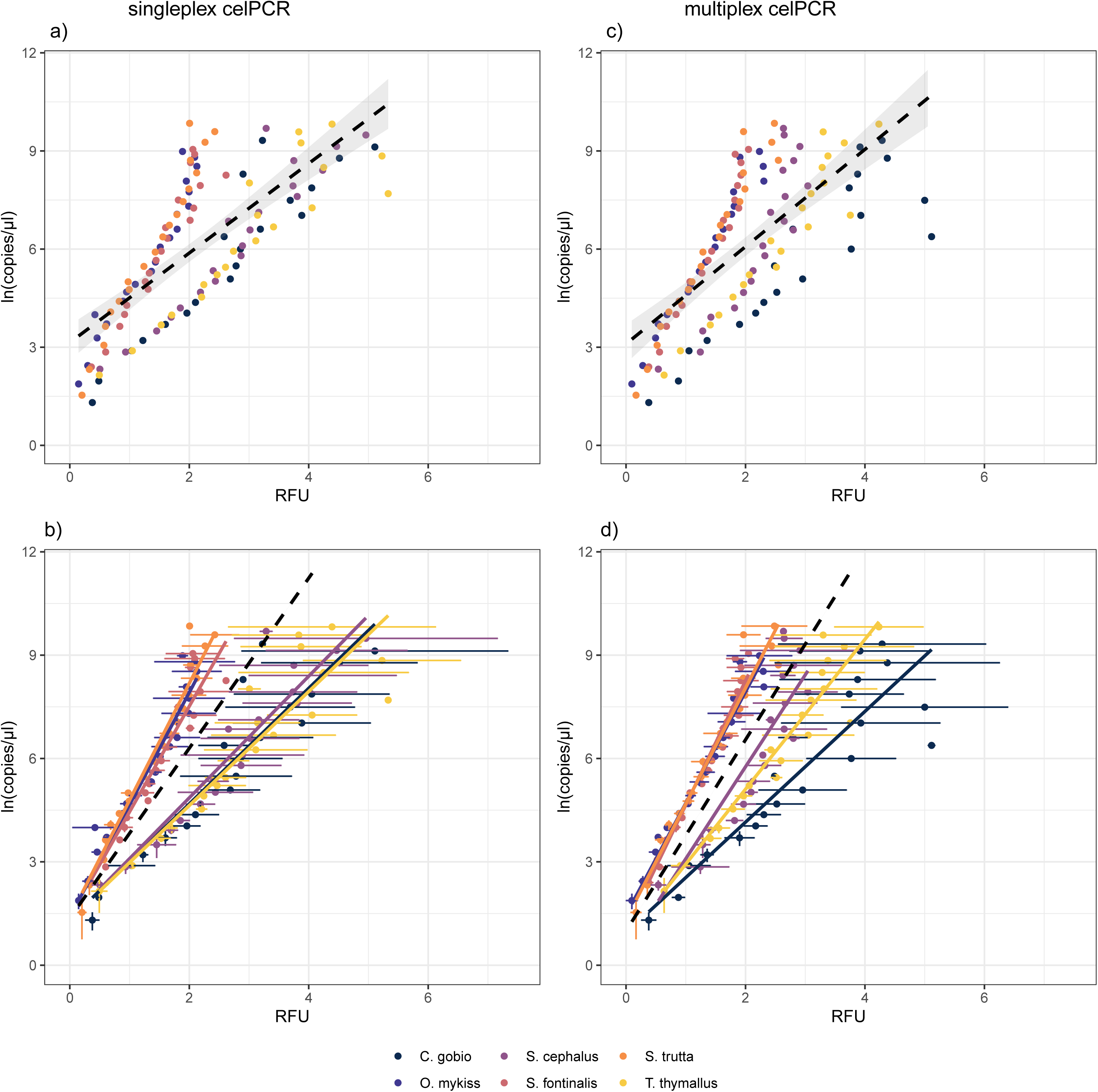
Linear models and mixed effect models for singleplex and multiplex celPCR in relation to copy numbers per µl extract (logarithmically scaled): panels a) and c) display mean RFU and copy numbery per dilution step and color coded by species, the black dashed line represents a linear model fitted onto this dataset without taking into account target species identity, the shaded area depicts the 95%-CIs; see SI2 for model specifications. Panels b) and d) show the mixed effects model using target species identity as random effect and permitting random slope and intercept (Tab. 2). Dots represent mean RFU and copy numbers per dilution step, with the corresponding standard deviations displayed as whiskers. The black dashed line depicts the linear model of the population mean; colored lines are the slopes associated with the individual species.

The use of individually measured RFU to predicted copy numbers from the linear mixed effects models showed similar trends for both singleplex and multiplex celPCRs: at low target DNA levels, predicted copy numbers were higher than the originally measured copy numbers, which is visualized by the linear regression line and its 95%-CI above the 45°-line (Fig. 4). At higher target DNA levels, this trend was reversed. However, for *S. fontinalis* and *C. gobio* in singleplex celPCR and *S. cephalus* in multiplex celPCR, the 95%-CI does not include the 45°-line at both the lower and upper end of the investigated DNA concentrations.

**Figure 4:**
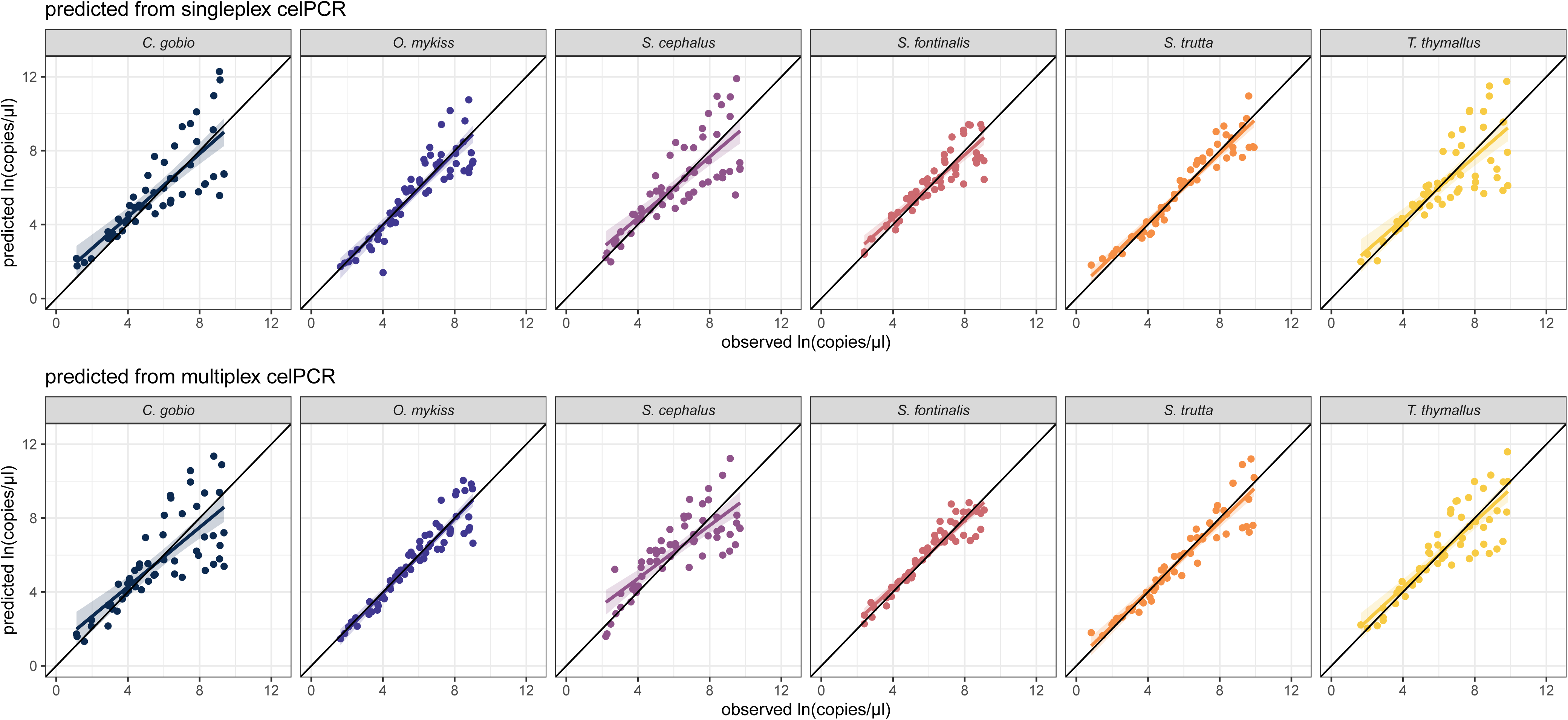
Observed copy numbers per µl extract (x-axis) plotted against the copy numbers predicted from the RFU obtained from singleplex celPCR (upper panel) and multiplex celPCR (lower panel). Per target species and PCR type, the comparisons are based on individual RFU obtained during the dilution series experiment. The black line (origin 0/0, slope 1) represents a perfect fit between observed and predicted copy numbers. For details on the linear models and 95% CIs illustrating the fit between observed and fitted copy numbers see SI2c.

The highest dilutions which produced positive amplifications (≥0.08 RFU; LOD) in singleplex and multiplex celPCRs contained target DNA quantities as measured via dPCR ranging from 0.6 to 8.1 copies per µl diluted extract. The LOQs in both singleplex and multiplex celPCRs inferred from triplicate dPCR measurements covered concentrations from 0.6 to 13 copies per µl extract (Table 4). As singleplex and multiplex PCRs both contained 1 µl of diluted extract, copies per µl are equivalent to copies in PCR.

**Table 4:** The limit of detection (LOD; lowest target DNA amount with amplification) and limit of quantification LOQ (lowest target DNA amount with all technical replicates yielding a positive result) of multiplex and singleplex celPCR for the different species

| species | LOD [copies/μl] | LOQ [copies/μl] |
| --- | --- | --- |
| *C. gobio* | 0.7 | 3.1 – 4.8 |
| *O. mykiss* | 6.5 – 8.1 | 5.1 – 8.0 (singleplex)<br>9.4 – 13 (multiplex) |
| *S. cephalus* | 0.6 – 2.4 | 9.1 – 12 (singleplex)<br>0.6 – 2.4 |
| *S. fontinalis* | 0.6 – 1.3 | 5.6 – 11 (singleplex)<br>0.6 – 1.3 |
| *S. trutta* | 0.6 (singleplex)<br>2.3 – 7.3 (multiplex) | 2.3 – 7.3 |
| *T. thymallus* | 1.8 – 2.4 | 5.2 – 13 |

Of the five field samples per target species which tested negative in multiplex celPCR, all but two were also negative in dPCR (one sample positive for *S. trutta* with 0.25 copies per µl extract and one positive for *C. gobio* with 0.13 copies per µl extract). The linear models describing for each primer pair the relationship between RFU and *ln*-transformed copy number in field-collected samples showed different R² levels ranging from 0.13 to 0.82 (SI2d, Fig. 5 upper panel). When comparing observed to predicted copy numbers, the data obtained from field samples fit well with data obtained from the dilution series experiment for *C. gobio*, *O. mykiss* and *S. trutta*. However, for all six primer pairs, the dispersion was higher for data derived from field-collected samples than for the dilution series data generated from tissue extracts (Fig. 5 lower panel). Ultimately, the observed and predicted copy numbers obtained from field-collected samples represent only a small part of the range examined via the dilution series and for *C. gobio*, *S. cephalus*, *S. fontinalis*, and *T. thymallus* align themselves at or beneath the lowest concentrations in the experiment (Fig. 5 lower panel).

**Figure 5:**
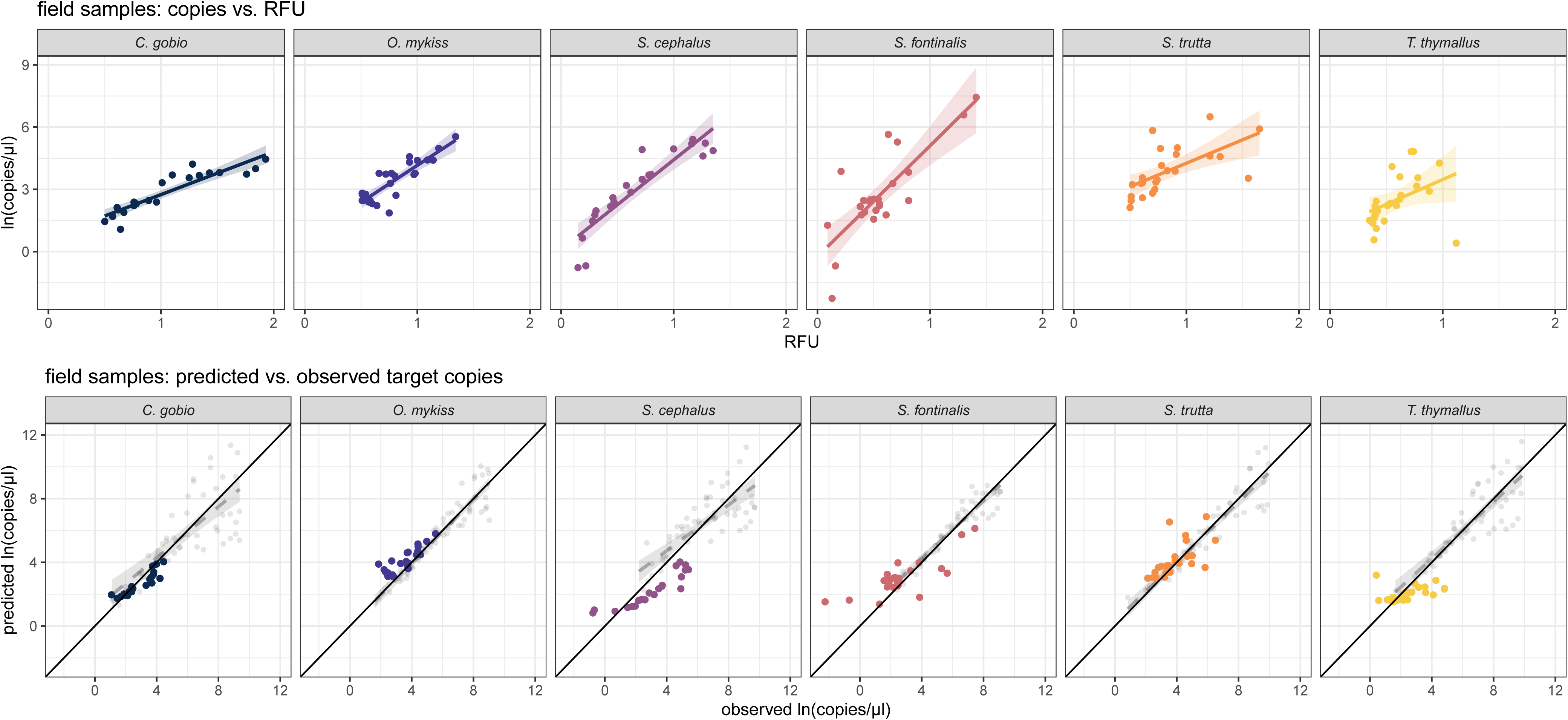
The relationship between RFU and *ln*-transformed target DNA copy numbers measured in field-collected samples for each of the primer pairs individually is displayed in the upper panel; for details on the linear models and their 95%-CIs see SI2d. The relationship between target DNA copy numbers measured in field-collected samples in comparison to the predicted copy numbers based on multiplex celPCR RFU of these field-collected samples is depicted in the lower panel. The black line (origin 0/0, slope 1) represents a perfect fit between observed and predicted copy numbers; the dashed regression line and the associated 95%-CIs are based on a comparison between measured and predicted copy numbers from the dilution series experiment. For details on the linear models and 95% CIs see SI2c.

## Discussion

Our results demonstrate the capacity of celPCR to provide a quantitative analysis of target eDNA copy number. After considering primer identity separately for singleplex and multiplex celPCR, it was possible to predict the number of target DNA copies in diluted extracts for each of the species-specific primer pairs. Furthermore, both singleplex and multiplex celPCR displayed high levels of sensitivity for diluted tissue extracts and field-collected eDNA samples, thus enabling the future application of cost-efficient multiplexes in large-scale screenings for target eDNA.

The comparison of DNA concentrations measured directly via dPCR to the signal strengths (RFU) measured via celPCR displays the exponential nature of endpoint PCR [19,52]. The diluted extracts processed simultaneously and in triplicate with both approaches showed increasing signal strength variability with increasing target DNA concentration. This is due to the endpoint reaction not being split into thousands of separate reactions [10]; hence, slight differences in DNA quantities at the start of the reaction can have strong effects on the final signal strengths. The signal strength in celPCR is also subject to saturation effects commonly occurring in the later stages of PCR and caused by template re-annealing, exhaustion of NTPs or primers, or loss of polymerase activity [53]. In our experiment, these two effects were visible for RFU > 3, and as none of the field-collected samples resulted in RFU > 2, they do not prevent the general semi-quantitative estimation of eDNA concentration from field-collected samples. Nevertheless, celPCRs of individual samples should be carried out in triplicate for accurate quantification, especially if higher target DNA concentrations are expected.

For the prediction of absolute target DNA concentrations from RFU it was necessary to account for primer effects, albeit the primer pairs were designed for equal amplification efficiency at uniform PCR conditions. As previously recommended [17,34], melting temperatures were as close as possible to 60 °C and the variation in fragment length (89-226 bp) was kept as small as possible and within the general suggestion for the detection of low concentrations of potentially degraded DNA from mixed samples [32]. The selected primers displayed minimal secondary structures and no competition for priming sites [17,34], and the multiplex PCRs were calibrated for equal amplification efficiency by adjusting primer concentrations in tests with target DNA templates [17,27]. However, all these measures were not sufficient to completely eliminate primer bias *a priori* for both singleplex and multiplex PCRs. A direct estimate of target DNA concentration was made possible by relating the RFU to absolute concentrations measured via dPCR and accounting for primer effects in linear mixed effects models. In our dilution series experiment, copy numbers predicted from singleplex or multiplex celPCR did not differ significantly from copy numbers measured with dPCR for most of the target species (exception *S. fontinalis*, *C. gobio* singleplex celPCR and *S. cephalus* multiplex celPCR) and the majority of individual copy numbers inferred from RFU fell inside the 95%-CI for concentrations below ~200 copies per µl extract. Therefore, absolute DNA concentrations can be deduced from RFU, if the efficiency of the applied primer pair(s) is directly compared between celPCR and a PCR-type enabling absolute quantification (i.e. dPCR). The resulting model permits predictions of the investigated target DNA concentration range. The amplification efficiency of a specific primer pair can differ between singleplex and multiplex PCRs despite careful design: For example, at equal primer concentrations, a less efficient primer pair leads to lower RFU in singleplex celPCR. Contrastingly, the concentration of a highly efficient primer pair needs to be reduced in multiplex celPCR to obtain comparable amplification success between all targets. Hence, it is necessary for quantitative estimations based on celPCR to evaluate the exact assay, which is going to be deployed in large-scale screenings, using qPCR or dPCR.

Both singleplex and multiplex celPCRs displayed similar levels of sensitivity in our experiments and resulted in positive amplifications of all reaction triplicates at concentrations between two and 13 target copies per µl extract (equalling two to 13 copies per 10 µl reaction volume). Depending on the target species, this was achieved at the highest or the second highest dilution step, where one or five target copies per µl extract were expected, respectively. At these low concentrations the stochastic nature of PCR causes some variation in detection success [19] and based on the number of replicates and the orders of magnitude covered in the dilution-series experiment, it was not possible to further refine the LODs and LOQs for each target species [22,23]. Nevertheless, our celPCRs showed sufficient sensitivity to detect target DNA in field-collected samples from rivers characterized by low productivity and low fish densities, and copy numbers in field-collected samples were predictable with the model obtained from the dilution series experiment, even though some signals were below the lower limit of the dilution series. If target DNA is expected to be present mostly at concentrations below 10 copies per µl extract (i.e. 10 copies per 10 µl PCR reaction volume), it is, however, possible to pre-amplify target DNA with a preceding singleplex PCR targeting for example all fish DNA contained in a sample [39]. Our results were consistent between dPCR, and multiplex celPCR, except for two field-collected samples, which tested negative in multiplex celPCR, but contained < 0.25 copies per µl extract in dPCR. Such low-concentration positives (below the LOD) have been previously observed in dPCR [22] and should be re-tested for further evaluation as these can be true positives, but also result from background signals of fluorescing foreign particles [54,55].

For all PCR platforms and visualization methods used in this study, a threshold is used to differentiate negative from positive results. In dPCR, this separates positive from negative droplets [10,56], whereas the lowest fluorescence signal distinctly different from background noise needs to be specified for both qPCR [22] and capillary electrophoresis [17]. The detection threshold of ≥ 0.08 RFU employed for both singleplex and multiplex celPCRs enabled the clear distinction of successful amplification from background fluorescence and was chosen based on previously used thresholds (0.07 and 0.1 RFU [27,57]) and after reviewing background signals in PCR and extraction negative controls. In dPCR, we chose to set a conservative threshold right below the cloud of positive droplets [56], therefore the amplitude of the threshold varied depending on the length of the target fragment. The use of EvaGreen Supermix made results directly comparable between celPCR and dPCR since the same primer pairs were used. However, this dPCR chemistry should be used with care, as the levels of background fluorescence can vary between field-collected samples.

The possibility for quantification via multiplex celPCR is appealing for target eDNA detection from high sample numbers. Especially commercial providers of eDNA services and smaller laboratories, which do not always have access to the newest technological advances, could benefit from this sensitive and cost-efficient approach (Table 1) when handling large sample numbers. While the semi-quantitative assessment of eDNA levels contained in field-collected samples is possible via celPCR after designing specific primers and optimizing the celPCR for maximum sensitivity, direct inference of the DNA concentration in the sample and absolutely quantitative comparisons between target species are only possible when accounting for primer effects and calibrating celPCR results using dPCR. Nevertheless, multiplex celPCR is a highly sensitive and broadly applicable approach for the detection and quantification of eDNA and will enable efficient and large-scale screenings in the context of species distribution monitoring at more affordable costs.

## Acknowledgements

This research was funded by the Austria Research Promotion Agency (FFG; project number 853219 awarded to MT). We are very grateful to D. Kirschner for his support in the collection of water samples, and D. Sint, M. Böcker and the members of the “Steinke Lab” for helpful comments on earlier manuscript versions and for pointing out additional references.

## Author contribution statement

MT conceived the study; BT and MT were responsible for study design. BT and YP carried out field sampling and laboratory analyses, BT analysed the data and wrote the first draft of the manuscript which was revised by YP and MT.

## Data Availability Statement

All singleplex and multiplex celPCR and dPCR data obtained during the dilution series experiment and the analysis of field-collected samples have been uploaded to Figshare and are available at https://doi.org/10.6084/m9.figshare.13139645.v1

