## Supplementary Information 1 for "Endpoint PCR coupled with capillary electrophoresis (celPCR) provides sensitive and quantitative measures of environmental DNA in singleplex and multiplex reactions"

**Supplementary Information 2**

**Bettina Thalinger^1,2^, Yannick Pütz^2^ & Michael Traugott^2,3^**

^1^ Centre for Biodiversity Genomics, University of Guelph, 50 Stone Road E, N1G 2W1, Guelph, Ontario, Canada

^2^ Department of Zoology, University of Innsbruck, Technikerstr. 25, 6020, Innsbruck, Austria

³ Sinsoma GmbH, Lannes 6, 6176 Voels, Austria

***Corresponding author:**

Bettina Thalinger,

Centre for Biodiversity Genomics, University of Guelph, 50 Stone Road E, N1G 2W1, Guelph, Ontario, Canada

**SI2a:** Linear models with RFU from singleplex celPCR (SP RFU) as predictor for RFU from multiplex celPCR. Per species, models are based on the mean values per dilution step. Columns describe the target species, adjusted R², the predictor variable, their parameter estimates, standard errors 95% confidence intervals, t-value, and p-value.

| **species** | **R² adj.** | **predictor variable** | **parameter estimate** | **SE** | **lower**  **95% CI** | **upper**  **95% CI** | **t-value** | **p-value** |  |
| --- | --- | --- | --- | --- | --- | --- | --- | --- | --- |
| *C. gobio* | 0.68 | intercept | 0.59 | 0.41 | -0.27 | 1.44 | 1.44 | 0.17 |  |
|  |  | SP RFU | 0.90 | 0.14 | 0.61 | 1.19 | 6.44 | <0.001 | *** |
| *O. mykiss* | 0.93 | intercept | 0.02 | 0.09 | -0.17 | 0.21 | 0.26 | 0.797 |  |
|  |  | SP RFU | 0.98 | 0.06 | 0.85 | 1.10 | 16.08 | <0.001 | *** |
| *S. cephalus* | 0.79 | intercept | 0.79 | 0.18 | 0.41 | 1.16 | 4.41 | <0.001 | *** |
|  |  | SP RFU | 0.51 | 0.06 | 0.38 | 0.63 | 8.63 | <0.001 | *** |
| *S. fontinalis* | 0.91 | intercept | 0.12 | 0.10 | -0.09 | 0.33 | 1.19 | 0.25 |  |
|  |  | SP RFU | 0.83 | 0.06 | 0.71 | 0.96 | 13.79 | <0.001 | *** |
| *S. trutta* | 0.92 | intercept | 0.01 | 0.11 | -0.22 | 0.25 | 0.13 | 0.898 |  |
|  |  | SP RFU | 1.01 | 0.07 | 0.86 | 1.16 | 14.04 | <0.001 | *** |
| *T. thymallus* | 0.73 | intercept | 0.62 | 0.30 | -0.001 | 1.24 | 2.10 | 0.05 |  |
|  |  | SP RFU | 0.65 | 0.09 | 0.46 | 0.84 | 7.23 | <0.001 | *** |

**SI2b:** Linear models with RFU as predictor for *ln*-transformed copy numbers per µl extract. Models for both singleplex and multiplex celPCR data were calculated using the mean values per dilution step and without including target species identity as categorical variable. Columns describe the source of the predicted values, the target species, adjusted R², the predictor variable, its parameter estimates, standard errors, 95% confidence intervals, t-value, and p-value.

| **species** | **R² adj.** | **predictor variable** | **parameter estimate** | **SE** | **lower**  **95% CI** | **upper**  **95% CI** | **t-value** | **p-value** |  |
| --- | --- | --- | --- | --- | --- | --- | --- | --- | --- |
| Singleplex PCR | 0.55 | intercept | 3.14 | 0.27 | 2.61 | 3.68 | 11.52 | <0.001 | *** |
|  |  | RFU | 1.37 | 0.11 | 1.15 | 1.59 | 12.24 | <0.001 | *** |
| Multiplex PCR | 0.51 | intercept | 3.10 | 0.30 | 2.51 | 3.70 | 10.31 | <0.001 | *** |
|  |  | RFU | 1.49 | 0.13 | 1.22 | 1.75 | 11.11 | <0.001 | *** |

**SI2c:** Per primer pair and respective target species, the linear models describing the relationship between observed copies per µl extract and predicted copies per µl extract based on data from the dilution series experiment are displayed. Models were calculated separately with predictions from singleplex celPCR and multiplex celPCR RFU. Columns describe the source of the predicted values, the target species, adjusted R², the predictor variable (always the observed *ln*-transformed copy numbers), their parameter estimates, standard errors, 95% confidence intervals, t-value, and p-value.

| **Predicted from** | **species** | **R² adj.** | **predictor variable** | **parameter estimate** | **SE** | **lower**  **95% CI** | **upper**  **95% CI** | **t-value** | **p-value** |  |
| --- | --- | --- | --- | --- | --- | --- | --- | --- | --- | --- |
| SP | *C. gobio* | 0.66 | intercept | 1.00 | 0.53 | -0.06 | 2.06 | 1.89 | 0.06 |  |
|  |  |  | ln(copies / µl) | 0.85 | 0.09 | 0.68 | 1.03 | 9.92 | <0.001 | *** |
| SP | *O. mykiss* | 0.82 | intercept | 0.04 | 0.37 | -0.70 | 0.78 | 0.11 | 0.91 |  |
|  |  |  | ln(copies / µl) | 0.95 | 0.060 | 0.86 | 1.10 | 16.23 | <0.001 | *** |
| SP | *S. fontinalis* | 0.82 | intercept | 0.87 | 0.37 | 0.14 | 1.61 | 2.39 | 0.02 | * |
|  |  |  | ln(copies / µl) | 0.86 | 0.06 | 0.75 | 0.98 | 15.23 | <0.001 | *** |
| SP | *S. trutta* | 0.92 | intercept | 0.44 | 0.24 | -0.05 | 0.92 | 1.81 | 0.07 |  |
|  |  |  | ln(copies / µl) | 0.93 | 0.04 | 0.85 | 1.00 | 24.66 | <0.001 | *** |
| SP | *S. cephalus* | 0.64 | intercept | 0.86 | 0.57 | -0.27 | 1.99 | 1.52 | 0.13 |  |
|  |  |  | ln(copies / µl) | 0.87 | 0.08 | 0.70 | 1.04 | 10.21 | <0.001 | *** |
| SP | *T. thymallus* | 0.63 | intercept | 0.92 | 0.58 | -0.24 | 2.08 | 1.59 | 0.12 |  |
|  |  |  | ln(copies / µl) | 0.84 | 0.09 | 0.67 | 1.02 | 9.75 | <0.001 | *** |
| MP | *C. gobio* | 0.60 | intercept | 1.08 | 0.56 | -0.05 | 2.21 | 1.91 | 0.06 |  |
|  |  |  | ln(copies / µl) | 0.80 | 0.09 | 0.62 | 0.99 | 8.83 | <0.001 | *** |
| MP | *O. mykiss* | 0.89 | intercept | -0.08 | 0.28 | -0.64 | 0.49 | -0.28 | 0.78 |  |
|  |  |  | ln(copies / µl) | 1.00 | 0.05 | 0.91 | 1.10 | 21.92 | <0.001 | *** |
| MP | *S. fontinalis* | 0.90 | intercept | 0.62 | 0.26 | 0.09 | 1.14 | 2.37 | 0.02 | * |
|  |  |  | ln(copies / µl) | 0.91 | 0.04 | 0.82 | 0.99 | 22.15 | <0.001 | *** |
| MP | *S. trutta* | 0.89 | intercept | 0.30 | 0.29 | -0.28 | 0.88 | 1.04 | 0.30 |  |
|  |  |  | ln(copies / µl) | 0.94 | 0.04 | 0.85 | 1.02 | 20.85 | <0.001 | *** |
| MP | *S. cephalus* | 0.63 | intercept | 1.85 | 0.48 | 0.89 | 2.82 | 3.84 | <0.001 | *** |
|  |  |  | ln(copies / µl) | 0.72 | 0.07 | 0.57 | 0.87 | 9.85 | <0.001 | *** |
| MP | *T. thymallus* | 0.75 | intercept | 0.68 | 0.46 | -0.25 | 1.60 | 1.46 | 0.15 |  |
|  |  |  | ln(copies / µl) | 0.89 | 0.07 | 0.75 | 1.03 | 12.80 | <0.001 | *** |

**SI2d:** Per primer pair and respective target species, the field-sample-based linear models describing the relationship between multiplex celPCR RFU and *ln*-transformed copies per µl extract are displayed. Columns describe the source of the predicted values, the target species, adjusted R², the predictor variable, its parameter estimates, standard errors, 95% confidence intervals, t-value, and p-value.

| **species** | **R² adj.** | **predictor variable** | **parameter estimate** | **SE** | **lower**  **95% CI** | **upper**  **95% CI** | **t-value** | **p-value** |  |
| --- | --- | --- | --- | --- | --- | --- | --- | --- | --- |
| *C. gobio* | 0.82 | intercept | 0.69 | 0.25 | 0.17 | 1.21 | 2.76 | 0.01 | * |
|  |  | MP RFU | 2.06 | 0.22 | 1.61 | 2.52 | 9.47 | <0.001 | *** |
| *O. mykiss* | 0.74 | intercept | 0.60 | 0.36 | -0.14 | 1.34 | 1.69 | 0.11 |  |
|  |  | MP RFU | 3.56 | 0.43 | 2.67 | 4.45 | 8.26 | <0.001 | *** |
| *S. cephalus* | 0.82 | intercept | 0.09 | 0.35 | -0.65 | 0.84 | 0.26 | 0.80 |  |
|  |  | MP RFU | 4.33 | 0.46 | 3.38 | 5.28 | 9.51 | <0.001 | *** |
| *S. fontinalis* | 0.63 | intercept | -0.25 | 0.53 | -1.34 | 0.84 | -0.47 | 0.65 |  |
|  |  | MP RFU | 5.35 | 0.84 | 3.61 | 7.08 | 6.39 | <0.001 | *** |
| *S. trutta* | 0.37 | intercept | 2.00 | 0.53 | 0.90 | 3.10 | 3.75 | 0.001 | ** |
|  |  | MP RFU | 2.26 | 0.60 | 1.02 | 3.49 | 3.80 | <0.001 | *** |
| *T. thymallus* | 0.13 | intercept | 1.09 | 0.72 | -0.40 | 2.57 | 1.52 | 0.14 |  |
|  |  | MP RFU | 2.39 | 1.15 | -0.004 | 4.78 | 2.07 | 0.05 |  |
